## Supplementary figures and images for "From paleness to albinism: Contribution of *OCA2* exon 10 skipping to hypopigmentation"

### Fig S1

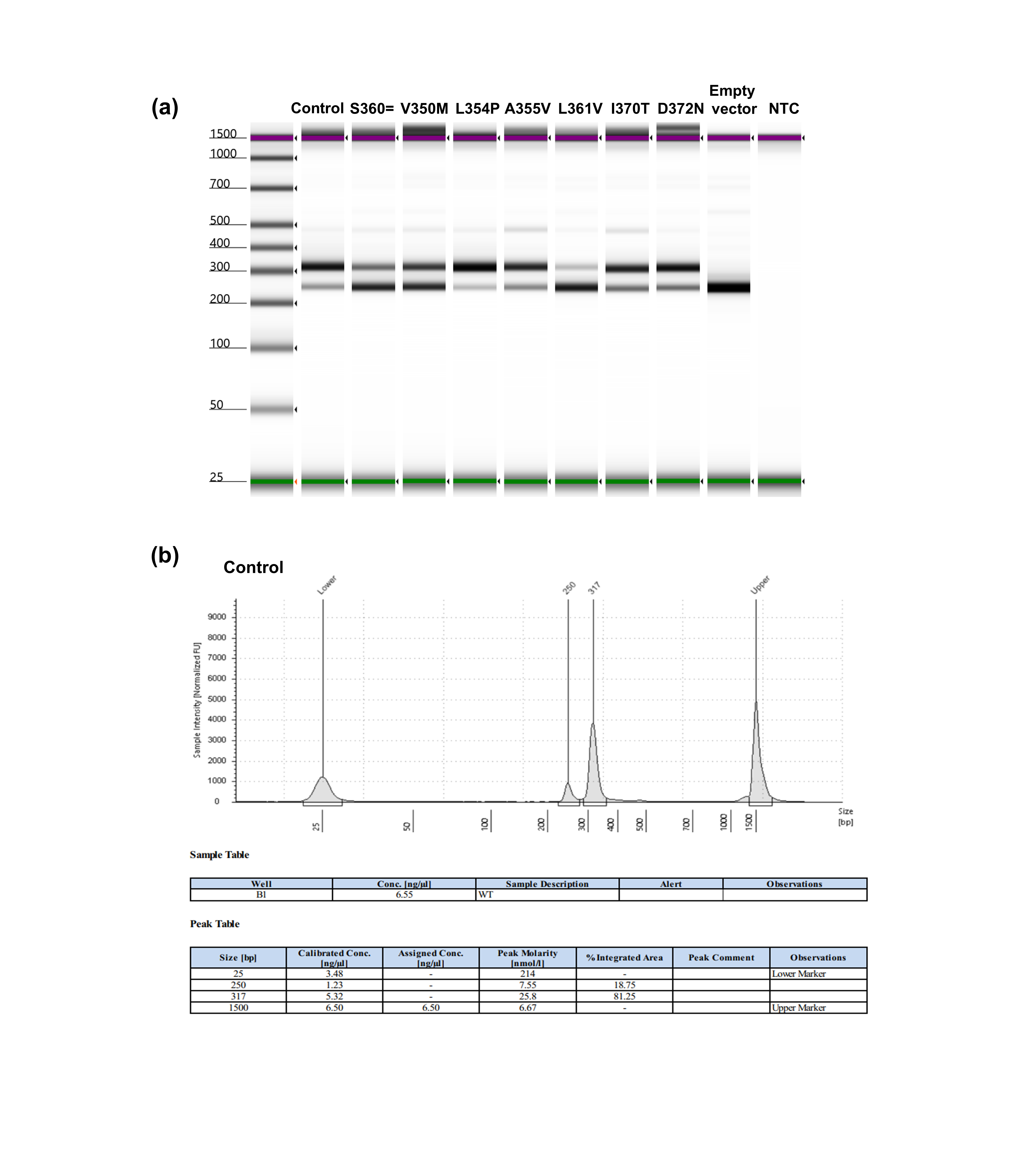

### Fig S2

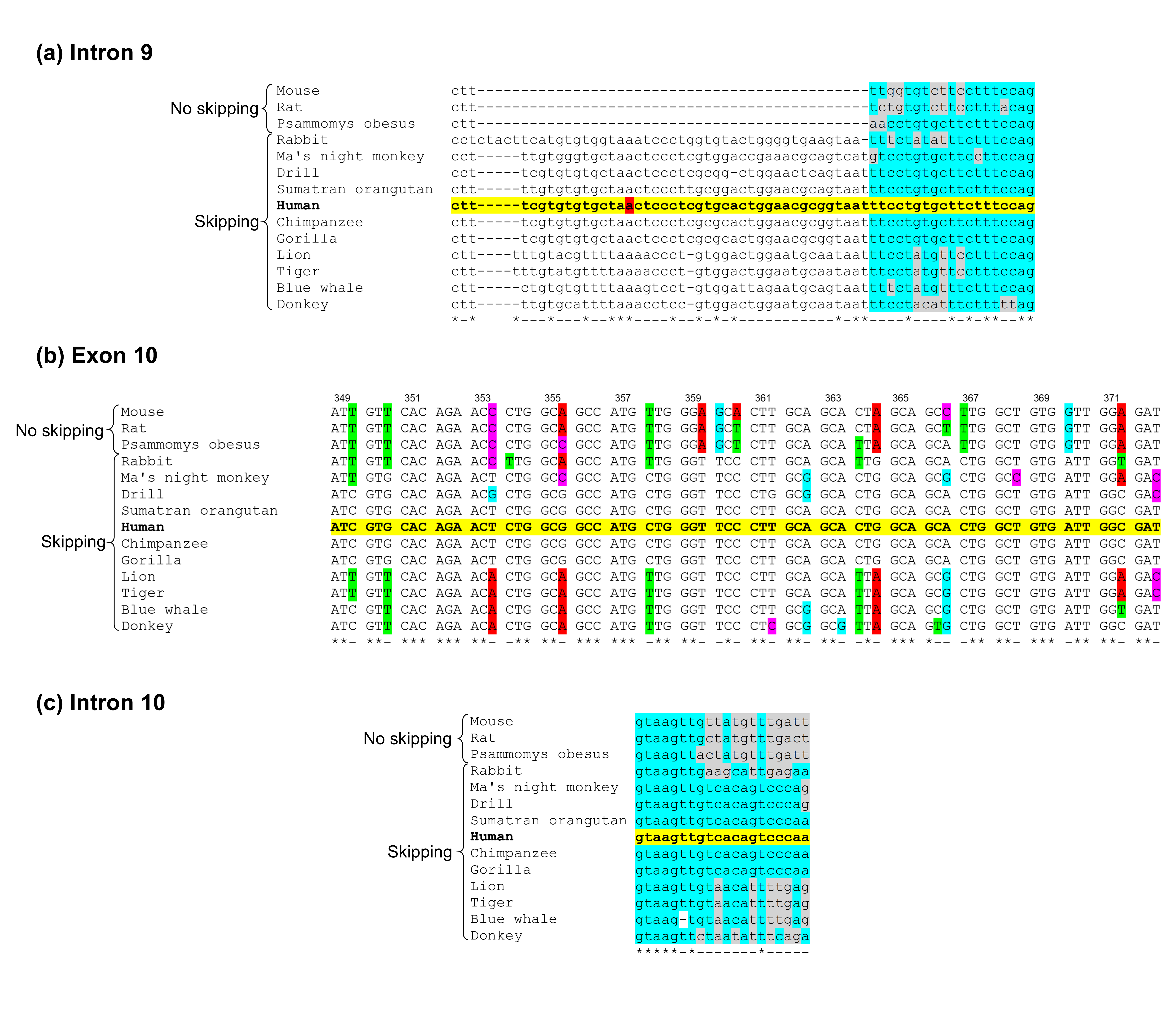

### Fig S3

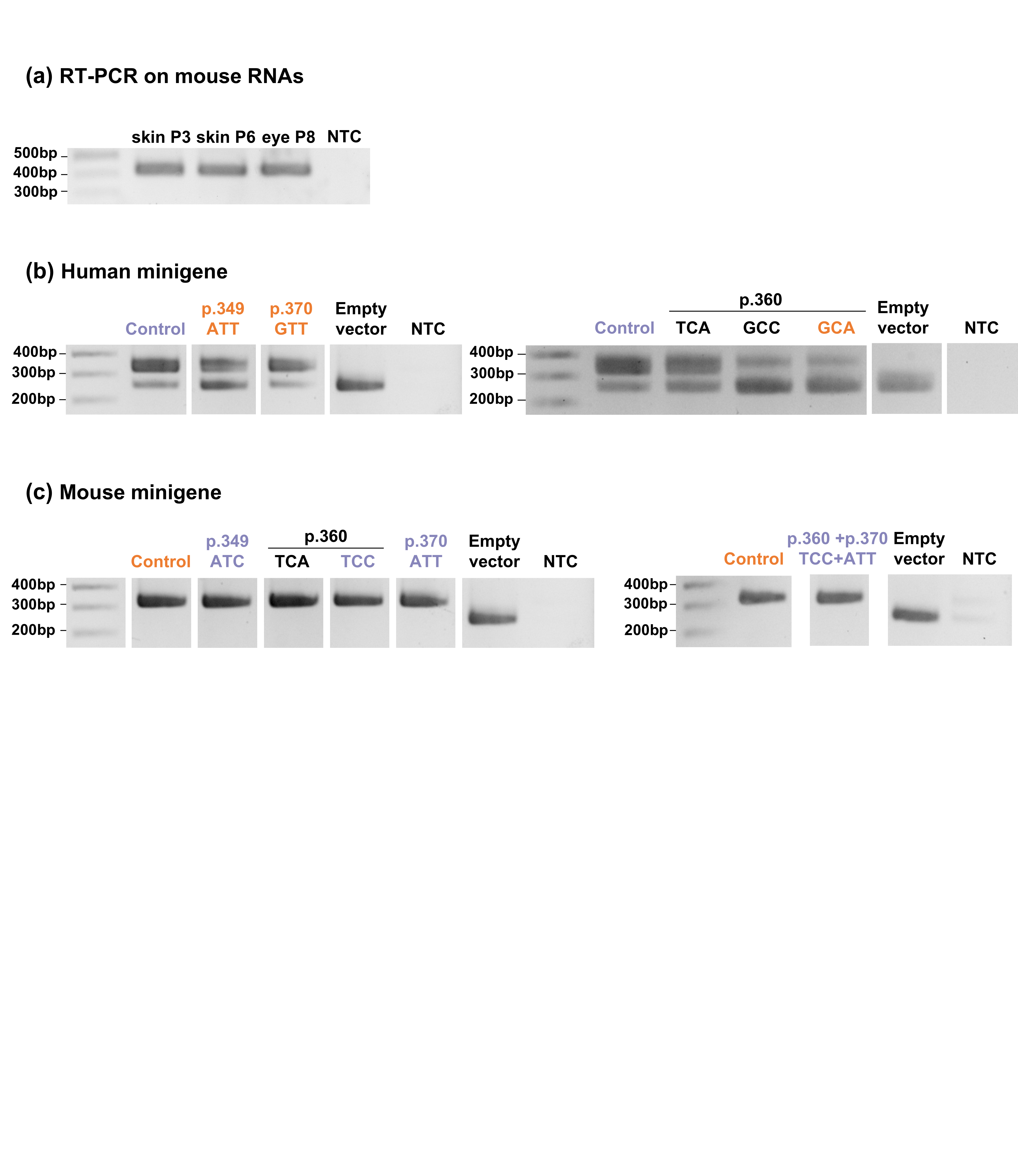

### Fig S4

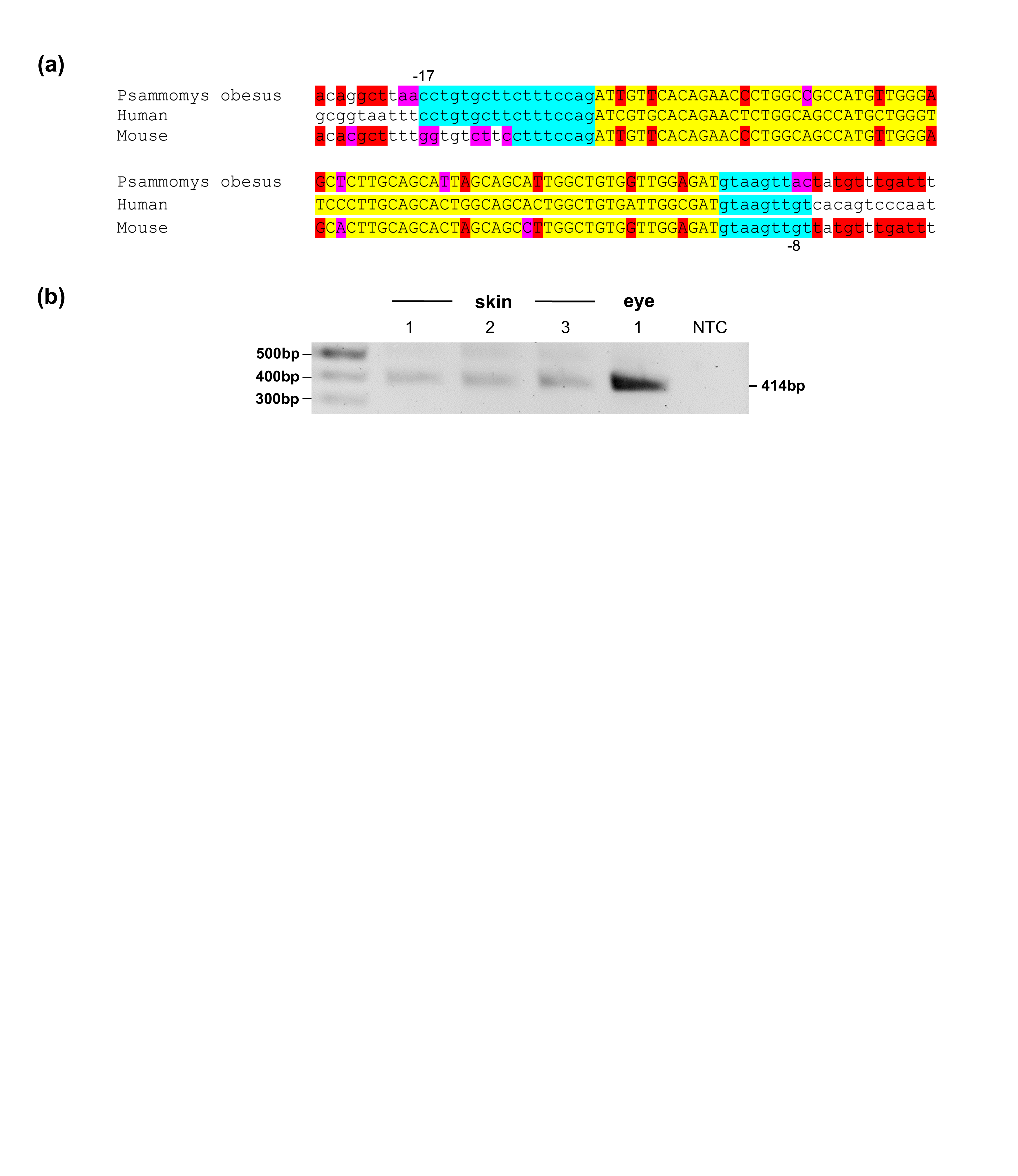

### Fig S5

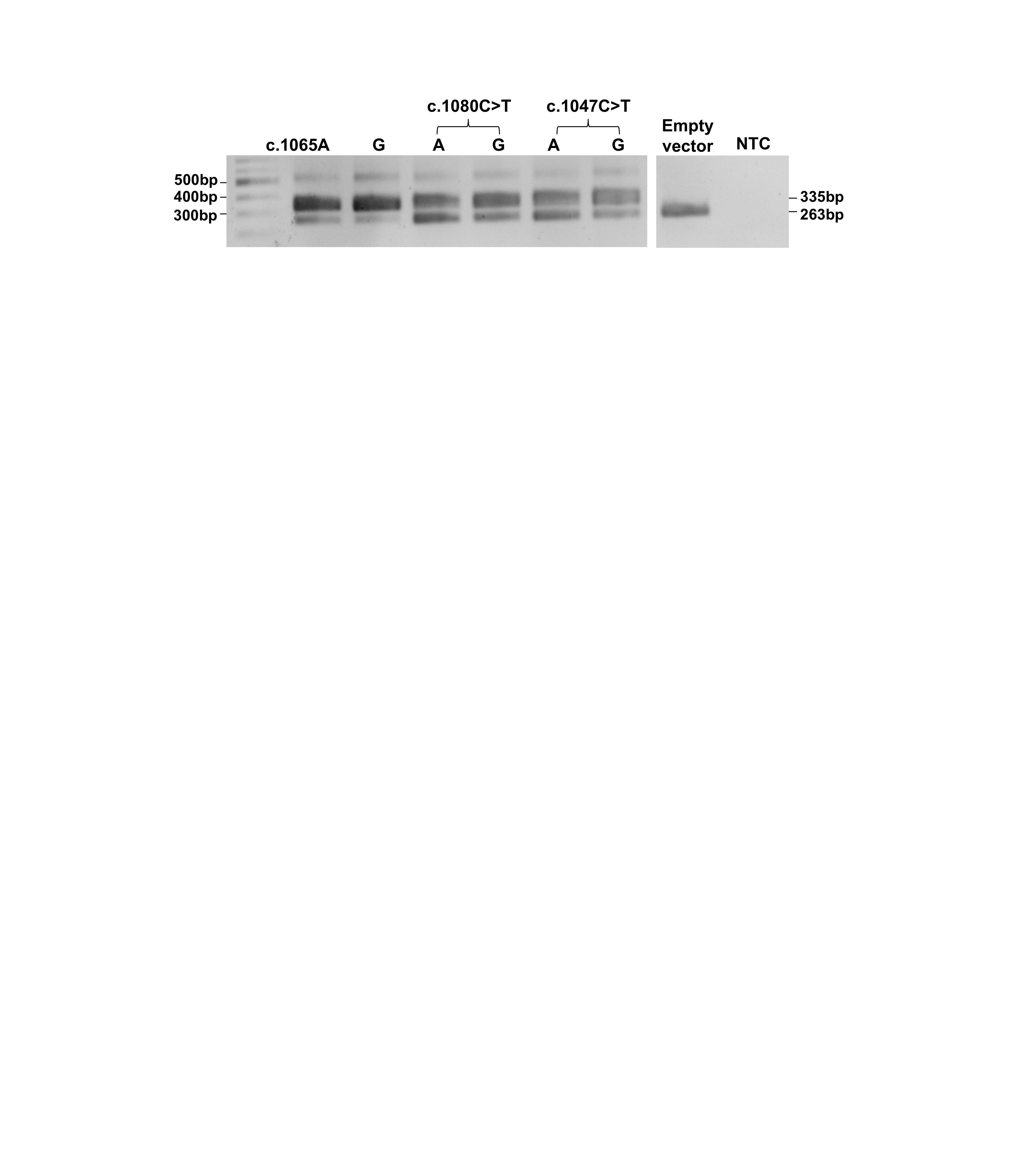

### Fig S6

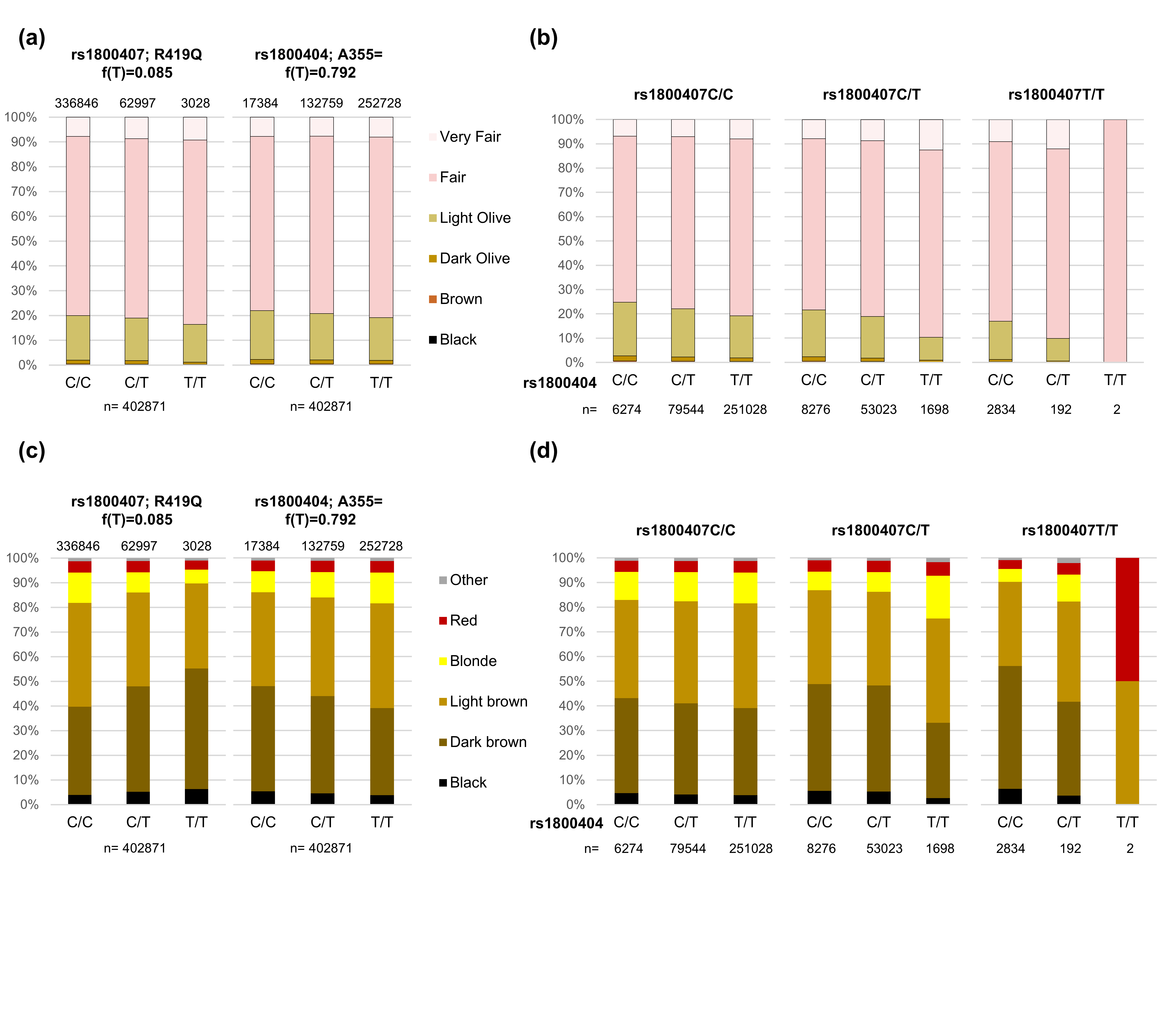

### Fig S7

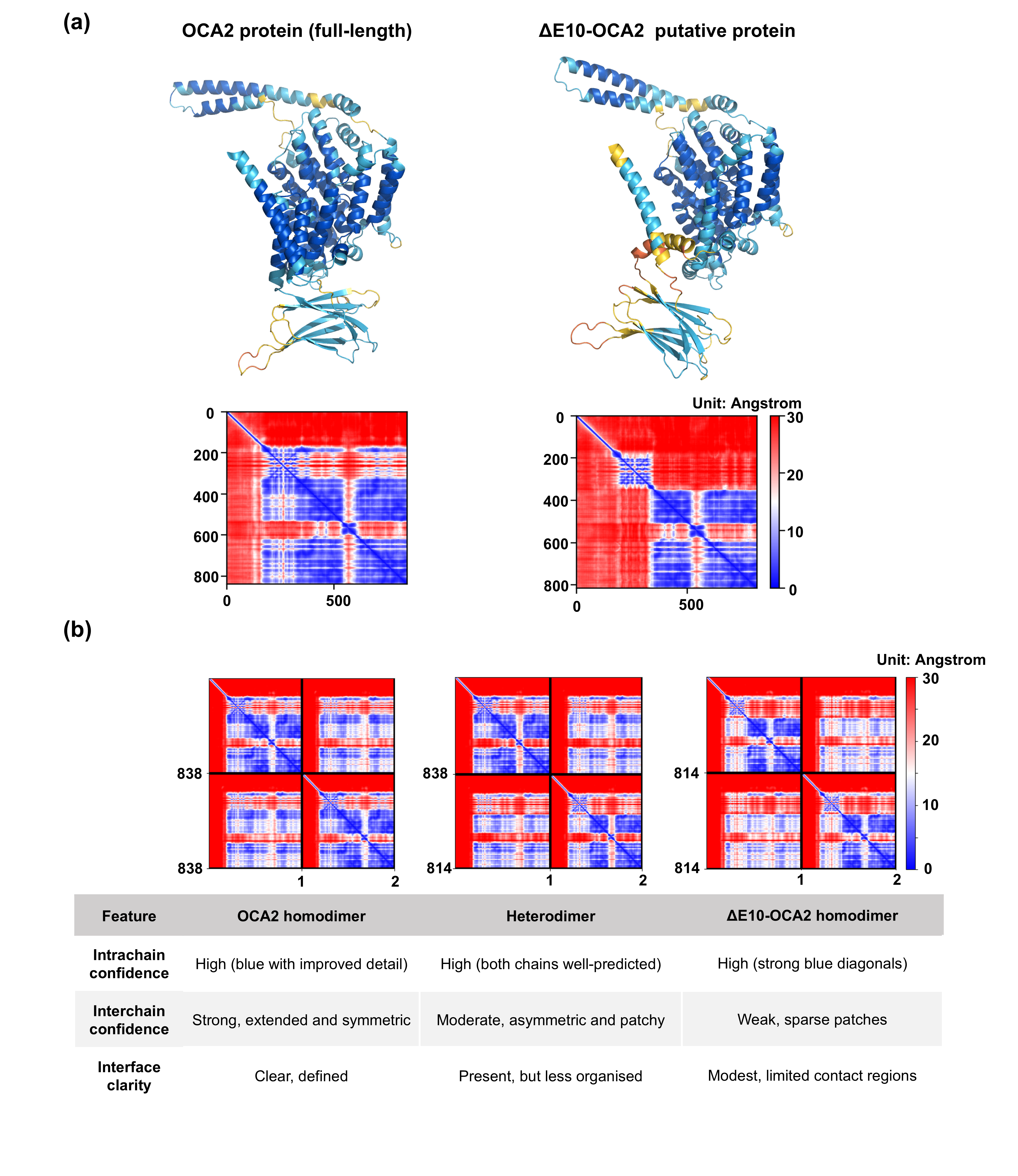

### Fig S8

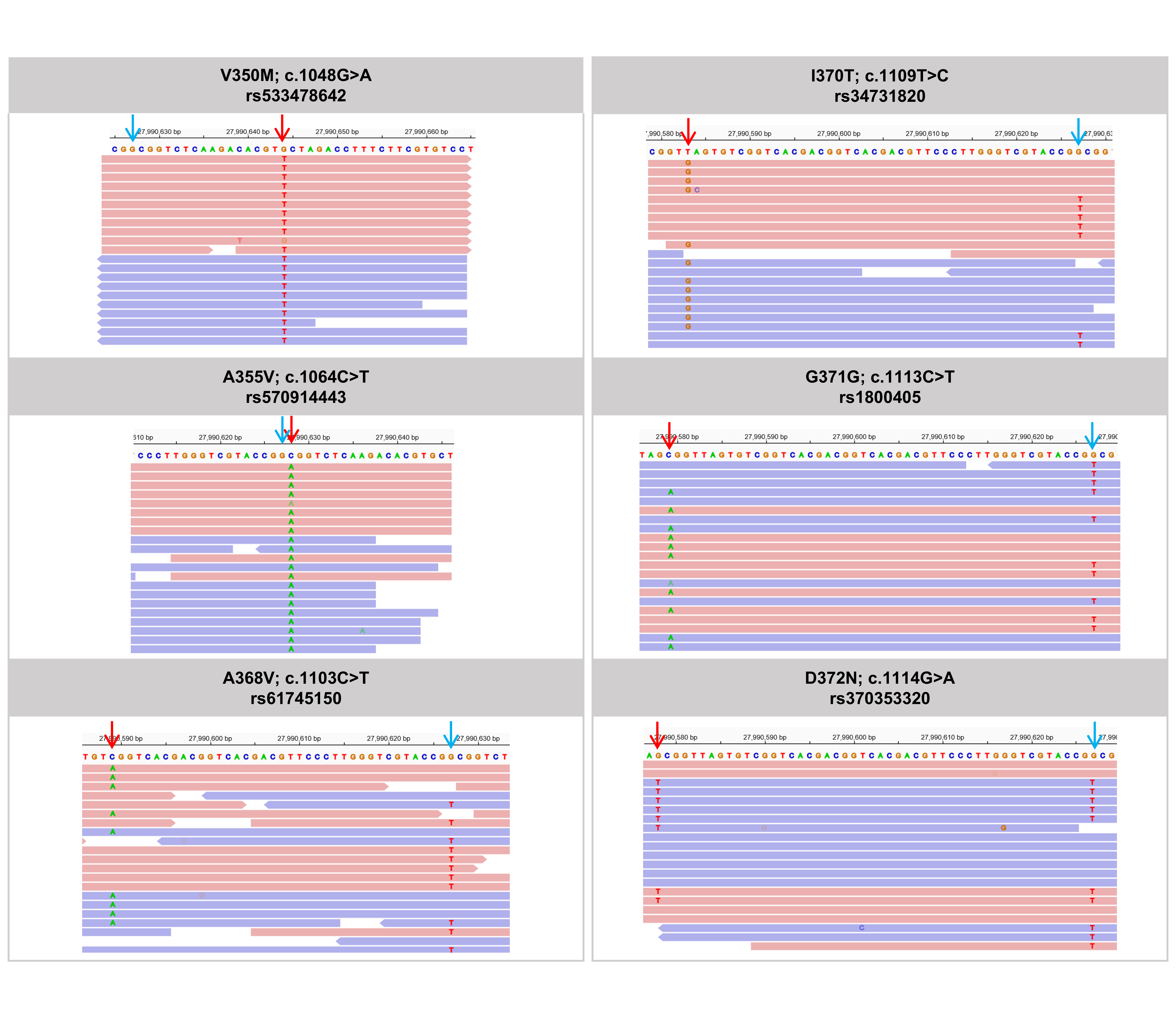
