## Supplementary material for "From paleness to albinism: Contribution of *OCA2* exon 10 skipping to hypopigmentation": Table S1

| Human minigene | | |
|---|---|---|
| Homologous recombination in pSPL3B vector | | Forward : 5’ ACCAGAATTCTGGAGCTCGAGCAGTTGCCAAGGGAGCATA 3’ |
| | | Reverse : 5’ ATCCTGCAGCGGCCGCTCGAGCCAAACAGGACACCCTCATC 3’ |
| Mutagenesis | c.1045-9T>G | Forward : 5’ GGTAATTTCCTGTGCTGCTTTCCAGATCGTGCAC 3’ |
| | | Reverse : 5’ GTGCACGATCTGGAAAGCAGCACAGGAAATTACC 3’ |
| | c.1047C>T;p.Ile349= | Forward : 5’ TGTGCTTCTTTCCAGATTGTGCACAGAACTCTGGC 3’ |
| | | Reverse : 5’ GCCAGAGTTCTGTGCACAATCTGGAAAGAAGCACA 3’ |
| | c.1048G>A ; p.Val350Met | Forward : 5’ GCTTCTTTCCAGATCATGCACAGAACTCTGG 3’ |
| | | Reverse : 5’ CCAGAGTTCTGTGCATGATCTGGAAAGAAGC 3’ |
| | c.1061T>C ; p.Leu354Pro | Forward : 5’ GTGCACAGAACTCCGGCAGCCATGCTG 3’ |
| | | Reverse : 5’ CAGCATGGCTGCCGGAGTTCTGTGCAC 3’ |
| | c.1064C>T ; p.Ala355Val | Forward : 5’ GCACAGAACTCTGGTAGCCATGCTGGGTTC 3’ |
| | | Reverse : 5’ GAACCCAGCATGGCTACCAGAGTTCTGTGC 3’ |
| | c.1065A>G ; p.Ala355= | Forward : 5’ TGCACAGAACTCTGGCGGCCATGCTGGGTTC 3’ |
| | | Reverse : 5’ GAACCCAGCATGGCCGCCAGAGTTCTGTGCA 3’ |
| | c.1078T>G ; p.Ser360Ala | Forward : 5’ GCCATGCTGGGTGCCCTTGCAGCAC 3’ |
| | | Reverse : 5’ GTGCTGCAAGGGCACCCAGCATGGC 3’ |
| | c.1080C>A ; p.Ser360= | Forward : 5’ GCCATGCTGGGTTCACTTGCAGCACTGGC 3’ |
| | | Reverse : 5’ GCCAGTGCTGCAAGTGAACCCAGCATGGC 3’ |
| | c.1078T>G + c.1080C>A | Forward : 5’ GCAGCCATGCTGGGTGCACTTGCAGCACTGGCA 3’ |
| | | Reverse : 5’ TGCCAGTGCTGCAAGTGCACCCAGCATGGCTGC 3’ |

| | | |
|---|---|---|
| | c.1081C>G ; p.Leu361Val | Forward : 5’ CATGCTGGGTTCCGTTGCAGCACTGG 3’ |
| | | Reverse : 5’ CCAGTGCTGCAACGGAACCCAGCATG 3’ |
| | c.1108A>G ; p.Ile370Val | Forward : 5’ GCAGCACTGGCTGTGGTTGGCGATGTAAGTT 3’ |
| | | Reverse : 5’ AACTTACATCGCCAACCACAGCCAGTGCTGC 3’ |
| | c.1109T>C ; p.Ile370Thr | Forward : 5’ CAGCACTGGCTGTGACTGGCGATGTAAGTTG 3’ |
| | | Reverse : 5’ CAACTTACATCGCCAGTCACAGCCAGTGCTG 3’ |
| | c.1114G>A ; p.Asp372Asn | Forward : 5’ CACTGGCTGTGATTGGCAATGTAAGTTGTCACAG 3’ |
| | | Reverse : 5’ CTGTGACAACTTACATTGCCAATCACAGCCAGTG 3’ |
| **Human minigene (intron 9: 73bp/ intron 10: 28bp)** 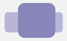 | | |
| Homologous recombination in pSPL3B vector | | Forward : 5’ GGAGCTCGAGCGGCCGCGTCCACACAGGCTTTCGTGTG 3’ |
| | | Reverse : 5’ GATCCTGCAGCGGCCGCGCCAGGGATTGGGACTGTG 3’ |
| **Human minigene (intron 9: 73bp/ intron 10: 85bp)** 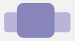 | | |
| Homologous recombination in pSPL3B vector | | Same forward as intron 9 73bp |
| | | Reverse : 5’ GATCCTGCAGCGGCCACCAGCGAAAGCCTGAATCC 3’ |
| **Human minigene (intron 9: 73bp/ intron 10: 212bp)** 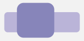 | | |
| Homologous recombination in pSPL3B vector | | Same forward as intron 9 73bp |
| | | Reverse : 5’ GATCCTGCAGCGGCCCCACTGGGATGTGAGTGTGTG 3’ |
| **Human minigene (intron 9: 73bp/ intron 10: 330bp)** 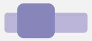 | | |
| Homologous recombination in pSPL3B vector | | Same forward as intron 9 73bp |
| | | Reverse : 5’ GATCCTGCAGCGGCCGCTAGGACGGTCCCCTCTAGTT 3’ |

| | | |
|---|---|---|
| **Human minigene (intron 9: 73bp/ intron 10: 429bp)** | | |
| Homologous recombination in pSPL3B vector | | Same forward as intron 9 73bp |
| | | Reverse : 5’ ATCCTGCAGCGGCCGCTCGAGCCAAACAGGACACCCTCATC 3’ |
| **Human minigene (intron 9: 326bp/ intron 10: 28bp)** | | |
| Homologous recombination in pSPL3B vector | | Forward : 5’ ACCAGAATTCTGGAGCTCGAGCAGTTGCCAAGGGAGCATA 3’ |
| | | Same reverse as intron 10 28bp |
| **Murine minigene** | | |
| Homologous recombination in pSPL3B vector | | Forward : 5’ GGAGCTCGAGCGGCCCTGAAGTCTCAATTAGATAAGGCAG 3’ |
| | | Reverse : 5’ GATCCTGCAGCGGCCTGGGTATCTCCAGGGCCTATG 3’ |
| Mutagenesis | c.1045-9C>G | Forward : 5’ CACGCTTTTGGTGTCTTGCTTTCCAGATTGTTCAC 3’ |
| | | Reverse : 5’ GTGAACAATCTGGAAAGCAAGACACCAAAAGCGTG 3’ |
| | c.1047T>C; p.Ile349= | Forward : 5’ GTCTTCCTTTCCAGATCGTTCACAGAACCCTGG 3’ |
| | | Reverse : 5’ CCAGGGTTCTGTGAACGATCTGGAAAGGAAGAC 3’ |
| | c.1078G>T; p.Ala360Ser | Forward : 5’ GCAGCCATGTTGGGATCACTTGCAGCACTAG 3’ |
| | | Reverse : 5’ CTAGTGCTGCAAGTGATCCCAACATGGCTGC 3’ |
| | c.1078G>T + c.1080A>C | Forward : 5’ GCAGCCATGTTGGGATCCCTTGCAGCACTAG 3’ |
| | | Reverse : 5’ CTAGTGCTGCAAGGGATCCCAACATGGCTGC 3’ |
| | c.1108G>A; p.Val370Ile | Forward : 5’ GCAGCCTTGGCTGTGATTGGAGATGTAAGTT 3’ |
| | | Reverse : 5’ AACTTACATCTCCAATCACAGCCAAGGCTGC 3’ |

| | | |
|---|---|---|
| **Murine minigene (intron 9: 75bp/ intron 10: 20bp)** 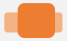 | | |
| Homologous recombination in pSPL3B vector | | Forward : 5’ GGAGCTCGAGCGGCCGCCTTTTTCATTTGATACACATGAAGC 3’ |
| | | Reverse : 5’ GATCCTGCAGCGGCCGCAATCAAACATAACAACTTACATCTC 3’ |
| **Minigene with mouse introns and human exon** 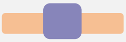 | | |
| Homologous recombination in pSPL3B vector | | Murine minigene forward and reverse primers |
| Mutagenesis | c.1045-9C>G | Forward : 5’ CGCTTTTGGTGTCTTGCTTTCCAGATCGTGC 3’ |
| | | Reverse : 5’ GCACGATCTGGAAAGCAAGACACCAAAAGCG 3’ |
| **Minigene with human introns and mouse exon** 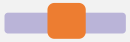 | | |
| Homologous recombination in pSPL3B vector | | Human minigene forward and reverse primers |
| Mutagenesis | c.1045-9T>G | Forward : 5’ CGCGGTAATTTCCTGTGCTGCTTTCCAGATTGTTC 3’ |
| | | Reverse : 5’ GAACAATCTGGAAAGCAGCACAGGAAATTACCGCG 3’ |
| **Minigene with human intron 9 and exon and mouse intron 10** 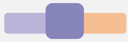 | | |
| Homologous recombination in pSPL3B vector | | Forward : 5’ GGAGCTCGAGCGGCCTACTCCATCTGGCCTTCC 3’ |
| | | Murine minigene reverse primer |
| **Minigene with mouse intron 9 and human exon and intron 10** 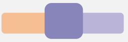 | | |
| Homologous recombination in pSPL3B vector | | Murine minigene forward primer |
| | | Human minigene reverse primer |

**Table S1:** List of primers for homologous recombination in pSPL3B and mutagenesis. Human exon and introns sequences are shown in dark and light purple respectively. Mouse exon and introns are in dark and light orange.
