## Supplementary material for "From paleness to albinism: Contribution of *OCA2* exon 10 skipping to hypopigmentation": Table S2

| Species | Gene identification | Exon 10 or equivalent | Full length *OCA2* transcript<br>Δ10 *OCA2* transcript | |
|---|---|---|---|---|
| Mouse | ENSMUSG00000030450 | No skipping | ENSMUST00000032633.12 Oca2-201 | https://www.ensembl.org/Mus_musculus/Gene/Summary?db=core;g=ENSMUSG00000030450;r=7:55889508-56186266 |
| Rat | ENSRNOG00000014465 | No skipping | ENSRNOT00000108070.2 Oca2-202 | https://www.ensembl.org/Rattus_norvegicus/Gene/Summary?db=core;g=ENSRNOG00000014465;r=1:116251796-116581838 |
| *Psammomys obesus* | 129676979 | No skipping | XM_055607524.1 | https://www.ncbi.nlm.nih.gov/gene/129676979 |
| Rabbit | ENSOCUG00000003510 | Skip | ENSOCUT00000003517.4 OCA2-201<br>ENSOCUT00000060813.1 OCA2-202 | https://www.ensembl.org/Oryctolagus_cuniculus/Gene/Summary?db=core;g=ENSOCUG00000003510;r=17:78036010-78397189 |
| Ma's night monkey | ENSANAG00000024732 | Skip | ENSANAT00000031480.1 OCA2-201<br>ENSANAT00000031481.1 OCA2-202 | https://www.ensembl.org/Aotus_nancymaae/Gene/Summary?db=core;g=ENSANAG00000024732;r=KZ202154.1:573255-828565 |
| Drill | ENSMLEG00000033698 | Skip | ENSMLET00000042501.1 OCA2-201<br>ENSMLET00000042507.1 OCA2-202 | https://www.ensembl.org/Mandrillus_leucophaeus/Gene/Summary?db=core;g=ENSMLEG00000033698;r=KN974395.1:2623632-2969172 |
| Sumatran ourangutan | ENSPPYG00000006278 | Skip | ENSPPYT00000007412.3 OCA2-201<br>ENSPPYT00000036919.1 OCA2-202 | https://www.ensembl.org/Pongo_abelii/Gene/Summary?db=core;g=ENSPPYG00000006278;r=15:2120028-2466892 |
| Human | ENSG00000104044 | Skip | ENST00000354638.8 OCA2-202<br>ENST00000353809.9 OCA2-201 | https://www.ensembl.org/Homo_sapiens/Gene/Summary?db=core;g=ENSG00000104044;r=15:27754875-28099315 |
| Chimpanzee | ENSPTRG00000006834 | Skip | ENSPTRT00000079363.1 OCA2-201<br>ENSPTRT00000045837.5 OCA2-202 | https://www.ensembl.org/Pan_troglodytes/Gene/Summary?db=core;g=ENSPTRG00000006834;r=15:8829677-9162797 |
| Gorilla | ENSGGOG00000005900 | Skip | ENSGGOT00000005933.3 OCA2-201<br>ENSGGOT00000030992.2 OCA2-202 | https://www.ensembl.org/Gorilla_gorilla/Gene/Summary?db=core;g=ENSGGOG00000005900;r=15:6233317-6581604 |
| Lion | ENSPLOG00000000481 | Skip | ENSPLOT00000000764.1 OCA2-201<br>ENSPLOT00000000775.1 OCA2-202 | https://www.ensembl.org/Panthera_leo/Gene/Summary?db=core;g=ENSPLOG00000000481;r=B4:26023193-26491686;t=ENSPLOT00000000764 |
| Tiger | ENSPTIG00000005972 | Skip | ENSPTIT00000006956.1 OCA2-201<br>ENSPTIT00000006959.1 OCA2-202 | https://www.ensembl.org/Panthera_tigris_altaica/Gene/Summary?db=core;g=ENSPTIG00000005972;r=KE722366.1:800302-1104135 |
| Blue whale | ENSBMSG00010005864 | Skip | ENSBMST00010008870.1 OCA2-201<br>ENSBMST00010008877.1 OCA2-202 | https://www.ensembl.org/Balaenoptera_musculus/Gene/Summary?db=core;g=ENSBMSG00010005864;r=7:2378692-2635994 |
| Donkey | ENSEASG00005019895 | Skip | ENSEAST00005082992.1 OCA2-205<br>ENSEAST00005031948.2 OCA2-203 | https://www.ensembl.org/Equus_asinus/Gene/Summary?db=core;g=ENSEASG00005019895;r=2:157758925-158119915 |

**Table S2:**
