## Supplementary material for "From paleness to albinism: Contribution of *OCA2* exon 10 skipping to hypopigmentation": Table S3

(a)

| rs1800404 | rs1800407 | Haplotype frequencies<br>Very Fair<br>N = 31 800 | Fair<br>N = 291 293 | Light Olive<br>N = 71 886 | Dark Olive<br>N = 6 189 | Brown<br>N = 1 680 | Black<br>N = 23 |
|---|---|---|---|---|---|---|---|
| T | C | 0.7925 | 0.7923 | 0.7781 | 0.7721 | 0.7842 | 0.7608 |
| T | T | 0.0042 | 0.0029 | 0.0014 | 0.0014 | 0.0003 | 0 |
| C | C | 0.1130 | 0.1218 | 0.1401 | 0.1509 | 0.1450 | 0.1522 |
| C | T | 0.090 | 0.0831 | 0.0804 | 0.0756 | 0.0705 | 0.0870 |
| **Additive haplotypic Odds Ratio** | | | | | | | |
| T | C | 1.077 [1.049 - 1.106] | ref | 0.853 [0.839 - 0.868] | 0.787 [0.748 - 0.827] | 0.831 [0.754 - 0.916] | 0.765 [0.438 - 1.336] |
| T | T | 1.569 [1.365 - 1.804] | ref | 0.437 [0.374 - 0.511] | 0.405 [0.243 - 0.674] | - | - |
| C | C | ref | ref | ref | ref | ref | ref |
| C | T | 1.169 [1.126 - 1.212] | ref | 0.840 [0.819 - 0.862] | 0.734 [0.678 - 0.795] | 0.716 [0.612 - 0.837] | 0.821 [0.640 - 1.052] |
| **Under the Assumption of proportionnal Odds** | | | | | | | |
| T | C | 0.853 [0.841 - 0.866] | $p = 2.05\ 10^{-101}$ | | | | |
| T | T | 0.472 [0.426 - 0.523] | $p = 2.56\ 10^{-46}$ | | | | |
| C | C | ref | | | | | |
| C | T | 0.813 [0.796 - 0.831] | $p = 1.10\ 10^{-78}$ | | | | |

(b)

| rs1800404 | rs1800407 | Haplotype frequencies<br>Black<br>N =16 679 | Dark Brown<br>N = 148 861 | Light Brown<br>N = 166 977 | Blonde<br>N = 46 666 |
|---|---|---|---|---|---|
| T | C | 0.76204 | 0.77222 | 0.79861 | 0.81809 |
| T | T | 0.00179 | 0.00228 | 0.00272 | 0.00385 |
| C | C | 0.12821 | 0.12735 | 0.12335 | 0.12305 |
| C | T | 0.10796 | 0.09815 | 0.07531 | 0.05502 |
| **Additive haplotypic Odds Ratio** | | | | | |
| T | C | 0.917 [0.886 - 0.949] | 0.936 [0.922 - 0.95] | ref | 1.027 [1.005 - 1.05] |
| T | T | 0.641 [0.483 - 0.852] | 0.807 [0.725 - 0.899] | ref | 1.419 [1.244 - 1.619] |
| C | C | ref | ref | ref | ref |
| C | T | 1.399 [1.333 - 1.468] | 1.263 [1.235 - 1.291] | ref | 0.732 [0.706 - 0.759] |
| **Under the Assumption of proportionnal Odds** | | | | | |
| T | C | 0.932 [0.92 - 0.944] | $p = 7.24\ 10^{-27}$ | | |
| T | T | 0.692 [0.633 - 0.756] | $p = 3.44\ 10^{-16}$ | | |
| C | C | ref | | | |
| C | T | 1.376 [1.35 - 1.402] | $p = 7.99\ 10^{-236}$ | | |

**Table S3**
