## Supplementary material for "From paleness to albinism: Contribution of *OCA2* exon 10 skipping to hypopigmentation": Table S4

| Variant | rs code | Publications | Allele frequency | Number of homozygotes | Frequency | | | | | | |
|---|---|---|---|---|---|---|---|---|---|---|---|
| | | | | | Europe | Africa | South Asia | East Asia | Middle East | Amixed americans | Rest of the world |
| p.Val350Met c.1048G>A | rs533478642 | PMID10649493 R.Kerr and al. 2000 PMID37882226 Beili Jiang and al. 2024 | 9.79E-05 | 2 | 2.50E-05 | | 1.30E-03 | 2.20E-05 | | | 1.40E-04 |
| p.Leu354Pro c.1061C>T | rs1008522653 | - | 6.82E-06 | 0 | | | | | | | |
| p.Ala355Ala c.1065G>A | rs1800404 | na | 7.17E-01 | 438294 | 8.10E-01 | 2.00E-01 | 3.90E-01 | 4.00E-01 | 6.70E-01 | 5.40E-01 | 6.80E-01 |
| p.Ala355Val c.1064C>T | rs570914443 | PMID23504663 Dimitre R Simeonov ans al. 2013 PMID28266639 Mohsin Shahzad ans al. 2017 PMID37882226 Beili Jiang and al. 2024 | 1.46E-04 | 4 | 1.20E-05 | 1.30E-05 | 2.20E-03 | 4.50E-05 | | 6.70E-05 | 2.10E-04 |
| p.Leu361Val c.1081C>G | rs769767739 | PMID37882226 Beili Jiang and al. 2024 | 6.20E-06 | 0 | | | | | | | |
| p.Ala368Val c.1103C>T | rs61745150 | PMID28451379 Jackson Gao and al. 2017 PMID32830442 Daniel Jackson and al. 2020 PMID37882226 Beili Jiang and al. 2024 | 1.32E-04 | 1 | 2.50E-06 | 2.00E-02 | | | | 5.00E-05 | 1.10E-04 |
| p.Ile370Thr c.1109T>C | rs34731820 | PMID10649493 R.Kerr and al. 2000 PMID32966289 Jenna E Rayner and al. 2020 PMID37882226 Beili Jiang and al. 2024 | 6.13E-04 | 9 | 5.00E-06 | 1.20E-02 | 5.00E-05 | | | 3.80E-04 | 7.20E-03 |
| p.Gly371= c.1113C>T | rs1800405 | PMID21541274 Markus N Preising and al. 2011 PMID37882226 Beili Jiang and al. 2024 | 1.40E-02 | 479 | 1.10E-02 | 3.00E-03 | 2.50E-02 | 1.60E-04 | 1.20E-02 | 9.80E-02 | 1.30E-02 |
| p.Asp372Asn c.1114G>A | rs370353320 | - | 1.92E-05 | 1 | 6.80E-06 | 9.30E-05 | 1.50E-04 | | | | 3.20E-05 |

**Table S4**
